# A Comparison of Methods for Estimating Substitution Rates from Ancient DNA Sequence Data

**DOI:** 10.1101/162529

**Authors:** K. Jun Tong, David A. Duchêne, Sebastián Duchêne, Jemma L. Geoghegan, Simon Y.W. Ho

## Abstract

The estimation of evolutionary rates from ancient DNA sequences can be negatively affected by among-lineage rate variation and non-random sampling. Using a simulation study, we compared the performance of three phylogenetic methods for inferring evolutionary rates from time-structured data sets: root-to-tip regression, least-squares dating, and Bayesian inference. Our results show that these methods produce reliable estimates when the substitution rate is high, rate variation is low, and samples of similar ages are not phylogenetically clustered. The interaction of these factors is particularly important for Bayesian estimation of evolutionary rates. We also inferred rates for time-structured mitogenomic data sets from six vertebrate species. Root-to-tip regression estimated a different rate from least-squares dating and Bayesian inference for mitogenomes from the horse, which has high levels of among-lineage rate variation. We recommend using multiple methods of inference and testing data for temporal signal, among-lineage rate variation, and phylo-temporal clustering.

## Introduction

Estimating the rate of molecular evolution is a key step in inferring evolutionary timescales and demographic dynamics from genetic data. Evolutionary rates can be estimated using time-structured genetic data sets, in which samples have been drawn at distinct points in time. In these cases, the molecular clock can be calibrated using the ages of ancient DNA sequences (Li et al. 1988; Rambaut 2000), as estimated by radiometric dating or stratigraphic correlation. Rates inferred from time-structured DNA data are essential to understanding evolutionary processes on short timescales (de Bruyn et al. 2011). Here, we examine some critical factors that can negatively affect these estimates of rates.

There are several different methods for estimating substitution rates from time-structured sequence data (Rieux & Balloux 2016). The simplest method is based on linear regression of root-to-tip (RTT) distances, measured in expected substitutions per site, against the ages of the corresponding sequences (Buonagurio et al. 1986). The slope of the regression line provides an estimate of the substitution rate. Two key drawbacks of this method are that the regression necessarily assumes a strict clock and that data points are not phylogenetically independent as some branches contribute to multiple root-to-tip measurements (Drummond et al. 2003; Rambaut et al. 2016). Least-squares dating (LSD) is another computationally efficient method that can estimate rates from time-structured data (To et al. 2016). It assumes a strict clock and fits a curve to the data using a normal approximation of the Langley-Fitch algorithm (Langley & Fitch 1974). This approximation is somewhat robust to departures from rate homogeneity among lineages. A third method is Bayesian phylogenetic analysis that can be used for joint estimation of substitution rates and the tree (Drummond et al. 2002). Bayesian methods can account for phylogenetic uncertainty and rate variation across branches, allow the error in sequence ages to be taken into account, and enable the co-estimation of evolutionary and demographic parameters of interest (Drummond et al. 2002). However, Bayesian analyses of time-structured sequence data have typically yielded very high rate estimates (Ho et al. 2007). Some of the statistical causes of high rates include tree imbalance (Duchêne et al. 2015a), closely related samples having the same age (Murray et al. 2015; Duchêne et al. 2015b), and extreme violations of the demographic assumptions (Navascués and Emerson 2009). The direction of bias is not always the same and depends on the pattern and magnitude of among-lineage rate variation (Wertheim et al. 2012).

As a particular form of time-structured data, ancient DNA presents unique analytical challenges compared to other time-structured data such as pathogen sequences. For instance, ancient sequences are typically scarce so that most of a data set is composed of contemporaneous sequences. The age of the root of ancient DNA studies is much older than in studies of pathogens, meaning that rates are more likely to vary between lineages. In this study, we examine two potential sources of error in rate estimates from time-structured sequence data: complex patterns of rate heterogeneity among lineages and sampling schemes in which samples with similar ages are also closely related. We investigate these factors in a simulation study and compare rate estimates made using three different methods from time-structured mitogenomic sequences.

## Results and Discussion

We simulated the evolution of DNA sequences under 12 different treatments, reflecting common evolutionary conditions many ancient DNA data sets. These treatments represented combinations of high (10^-7^ subs/site/year) and low (10^-8^ subs/site/year) substitution rates, three levels of among-lineage rate variation (high, medium, and low), and two levels of phylogenetic clustering (high and low). In general, the three methods of analysis (RTT regression, LSD, and Bayesian inference) produced more accurate estimates for sequence data produced by simulation using a high rate than with a low rate (Figure 1). The spread of estimates from each of the six high-rate treatments across all three methods was relatively narrow in most cases. The Bayesian median estimates were accurate, with a small spread, for all sequence data that had been produced with a high rate.

**FIG. 1.**
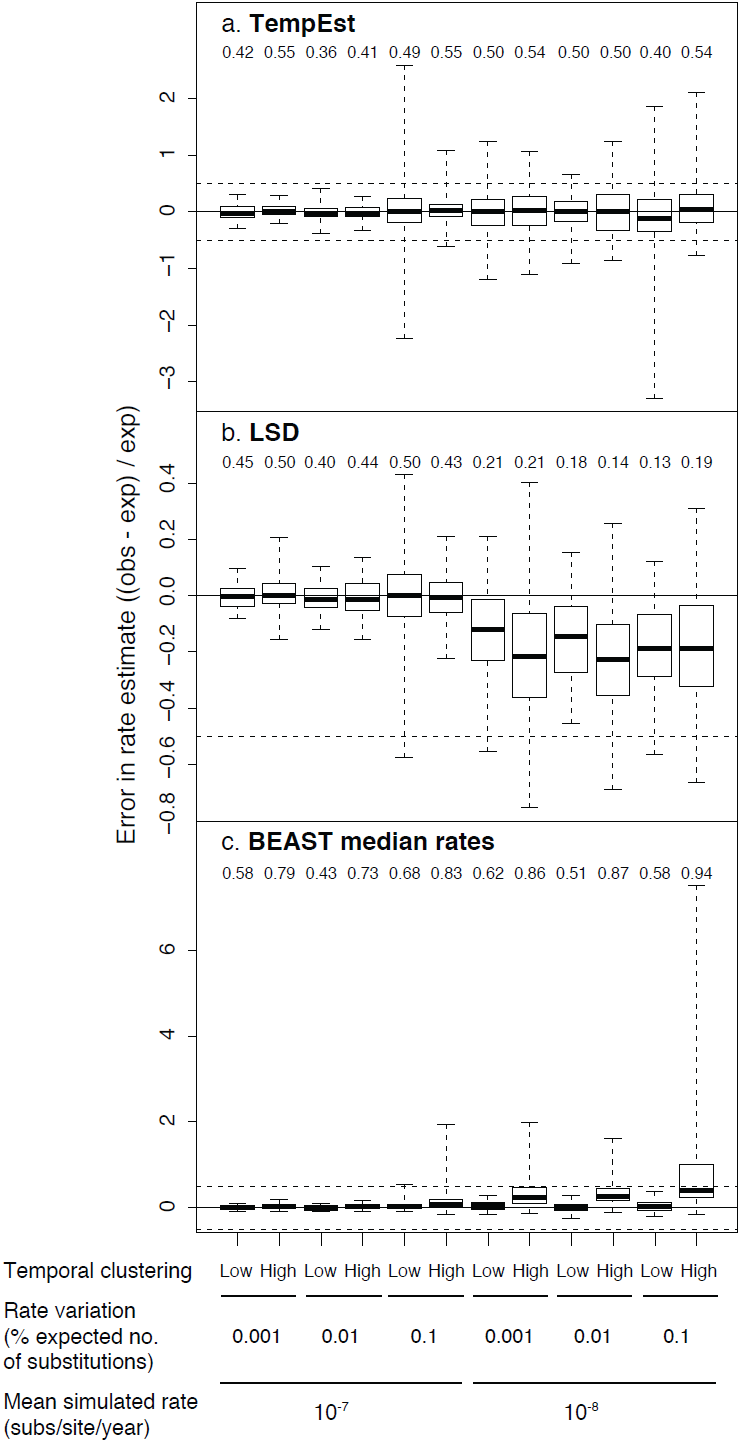
Estimates of substitution rates from sequence data produced under 12 different simulation conditions. Data were analysed using Bayesian inference in BEAST, least-squares dating in LSD, and root-to-tip regression in TempEst. Numbers on each panel indicate the proportion of estimates that are above the true simulated rate. Solid horizontal line indicates the true rate. Dashed horizontal lines indicate half a degree of magnitude above and below the true rate.

For sequence data simulated with a low rate, the mean estimates generally had a small spread except when there were high levels of phylogenetic clustering (see below). LSD mildly underestimated the rate for these data sets. RTT regression produced mean rate estimates with a greater spread for the data sets that had evolved with a low rate.

The presence of among-lineage rate variation increased the spread of mean rate estimates from sequence data that had evolved with a high rate. However, this rate variation did not have a measurable impact on the rate estimates made using RTT regression and Bayesian inference from the sequences that had evolved with a low rate.

Unexpectedly, high levels of phylogenetic clustering of sequences of similar ages were not associated with systematic over- or underestimation of the rate. Previous studies have shown that such phylogenetic clustering could obscure the temporal signal as assessed by the date randomization test (Duchêne et al. 2015b). We found that phylo-temporal clustering did not have much impact on the rate estimates from data sets that had evolved with a high rate, but tended to increase the spread of the mean estimates from the slowly evolving sequences. Notably, Bayesian inference was accurate and precise for data sets that had low levels of phylo-temporal clustering, regardless of whether they had evolved with a high or low rate.

The interaction between low rate, high among-lineage rate variation, and high phylo-temporal clustering is apparent in the Bayesian rate estimates (Figure 2a). This is consistent with the results of previous studies of these individual factors, including the effects of extreme rate variation among lineages (Wertheim et al. 2012) and phylo-temporal clustering (Duchêne et al. 2015b; Murray et al. 2015). RTT regression and LSD appear to be more robust to the interaction of these three unfavourable factors, although it is evident that a low rate, high among-lineage rate variation, and high phylo-temporal clustering all contribute to estimation error (Figure 1).

**FIG. 2.**
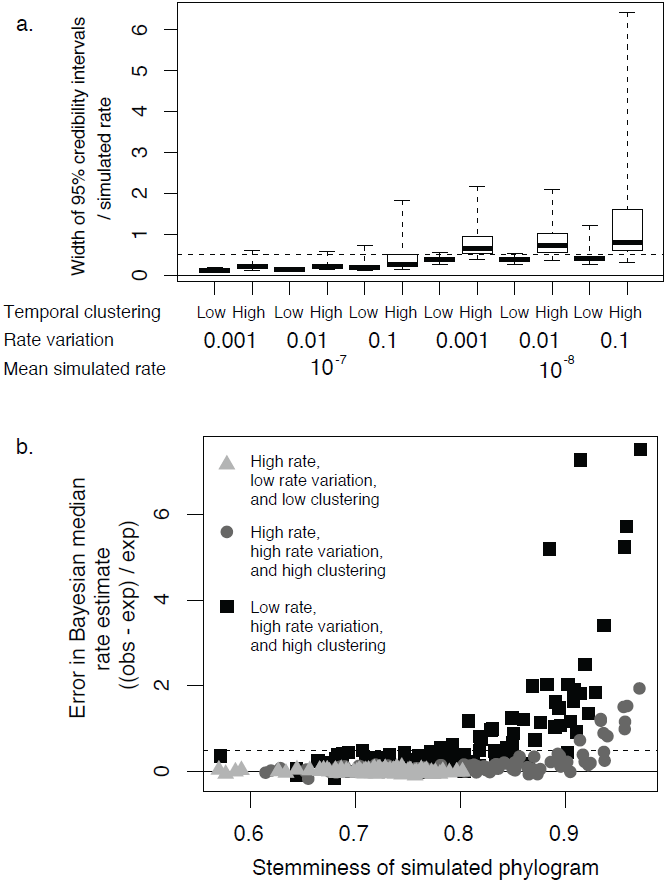
(*a*) Precision of Bayesian estimates of substitution rates across 12 simulation conditions, as measured by the width of the 95% credibility interval of the estimate divided by the rate used for simulation. One hundred data sets were produced by simulation under distinct evolutionary conditions and analysed using BEAST. (*b*) Relationship between phylogenetic stemminess and the error of rate estimate according to Bayesian inference. Stemminess corresponds to ratio of internal to terminal branch lengths.

The data sets that yielded erroneous rate estimates when analysed using Bayesian inference tended to have phylograms (trees with branch lengths proportional to genetic change) with a high ratio of internal to terminal branch lengths (Figure 2b). These trees are generally shorter than those with short internal and long terminal branches, such that there is less information from which to estimate the rate. We found a positive correlation between phylogenetic stemminess and the spread of median posterior rate estimates.

We also used the three methods to analyse six time-structured mitogenomic data sets, from Adélie penguin, brown bear, dog, horse, modern human, and woolly mammoth. All three methods produced rate estimates that were consistent with one another when the sequences had evolved according to a strict clock (Figure 3). Rate estimates were largely congruent with one another even when the data showed evidence of among-lineage rate variation, except for horse mitogenomes for which the rate estimate from RTT regression fell outside the 95% credibility interval of the Bayesian estimate.

**FIG. 3.**
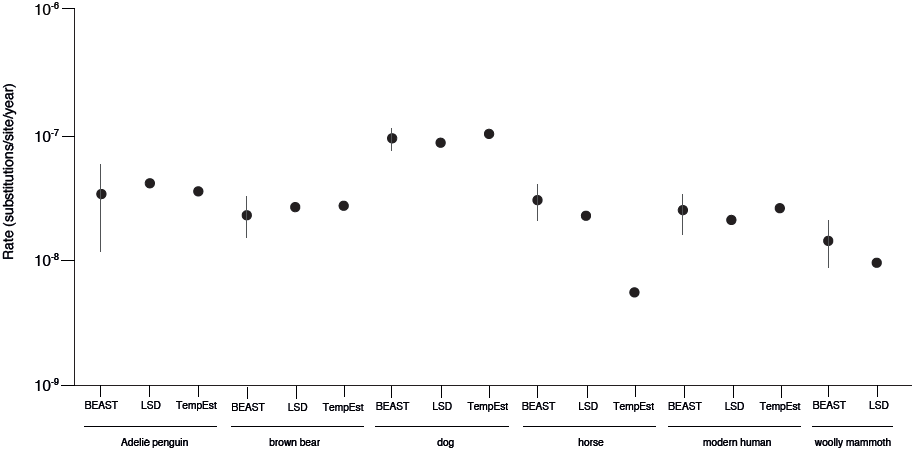
Estimates of substitution rates from six time-structured mitogenomic data sets. Bayesian estimates are indicated by their median and 95% credibility intervals. Main publications from which the sequence data were obtained: Adelié penguin, Subramanian et al. (2009); brown bear, Miller et al. (2012); dog, Thalmann et al. (2013); horse, Lippold et al. (2011), Achilli et al. (2012), and Orlando et al. (2013); modern human, Brotherton et al. (2013); and woolly mammoth, Gilbert et al. (2008). Details of the data sets are given in Supplementary Table S1.

## Conclusions

Our study has shown that three common methods of rate estimation from time-structured data produce robust estimates of substitution rates under most evolutionary conditions. Of the three methods, RTT regression is useful beyond its role as a ‘sanity check’: it produced rate estimates similar to those of LSD and Bayesian inference for four of the six mitogenomic data sets and performed well in our simulation study. RTT regression is also useful in informing model selection because it can be used to detect clocklike evolution, based on the fit to the data. Least-squares dating is particularly valuable for analyses of large data sets, for which the computational demands of a Bayesian phylogenetic analysis would be prohibitive (e.g. Mourad et al. 2015). It is remarkably robust to violations of the strict clock and can handle data with appreciable levels of among-lineage rate variation (To et al. 2016).

Our results also reveal that the three methods respond differently to the potentially confounding impacts of among-lineage rate variation and phylo-temporal clustering of sequences. This highlights the value of using all three methods to analyse time-structured sequence data. Increasing the reliability of rate estimates will lead to a more accurate understanding of demographic and evolutionary processes on recent timescales.

## Materials and Methods

### Simulations

We simulated genealogies of 100 tips in BEAST 2 (Bouckaert et al. 2014), under a constant population size and conditions that resemble ancient DNA studies of Pleistocene vertebrates. In all cases, the root height was fixed to 500,000 years and half of the tips corresponded to present-day samples. The ages of the other 50 tips were randomly distributed between the present and 50,000 years ago (i.e. 10% of the age of the root). Trees contained two scenarios of phylo-temporal clustering, with 100 replicates each: high clustering was simulated by making all present-day samples form a monophyletic group, whereas low clustering was simulated by only making half of the present-day samples form a monophyletic group.

Using the simulated genealogies and the program NELSI (Ho et al. 2015), we simulated the evolution of nucleotide sequences while varying the mean substitution rate and the degree of among-lineage rate variation under scenarios resembling time-structured mitogenomic data sets. Simulations were performed using two substitution rates that cover the range of rates in most molecular dating studies using ancient DNA: a high rate of 10^-7^ and low rate of 10^-8^ subs/site/year. For each of the two rate schemes, we simulated three scenarios of among-lineage rate variation under a white-noise model (Lepage et al. 2007), with variance along each branch of 0.1% (low), 1% (medium), and 10% (high) of the expected number of substitutions. Sequence evolution was simulated according to the HKY+Γ substitution model using the R package phangorn (Schliep et al. 2011) for each of the 100 tree replicates in the 12 different scenarios. All sequences had lengths of 15,000 nucleotides, to reflect the approximate size of many vertebrate mitogenomes.

For the 100 data sets in each simulation treatment, we used three methods to estimate the substitution rate: root-to-tip regression in TempEst 1.5 (Rambaut et al. 2016), least-squares fitting in LSD 0.3 (To et al. 2016), and Bayesian phylogenetic inference in BEAST 1.8.3 (Drummond et al. 2012). We performed a regression of root-to-tip distances against sample ages in TempEst for each data set. Substitution rates were also estimated in LSD, with the ages of the samples used to inform the least-squares fitting algorithm. We performed Bayesian phylogenetic analysis of each data set using BEAST. For the substitution rate, we used a continuous-time Markov chain reference prior (Ferreira & Suchard 2008).

### Mitochondrial Genomes

We used the three methods to analyse six time-structured mitogenomic data sets (Table S1). To infer phylograms for TempEst and LSD, we used maximum likelihood in RAxML 8.2.4 (Stamatakis 2014) with the HKY+Γ model of nucleotide substitution. In each case, a rapid bootstrapping analysis with 100 replicates was followed by a search for the best-scoring tree. Outgroup sequences were included in order to allow the position of the root to be estimated (see Table 1), but were pruned from the tree for subsequent analyses of substitution rates.

For each data set, we checked for temporal structure using a date-randomization test (Ramsden et al. 2009). In this test, the sample ages are randomly reassigned to the sequences and a rate is re-estimated. This is repeated several times to generate a set of rate estimates from date-randomized data sets. A data set is considered to have adequate temporal structure if its rate estimate differs from those obtained from the date-randomized data.

The data used here is available from https://github.com/kjuntong/aDNAratesproject

## Acknowledgements and funding information

This work was supported by the Australian Research Council (grant number FT160100167) and the University of Sydney HPC service. KJT was supported by an Australian Postgraduate Award. SD was supported by a McKenzie Fellowship from the University of Melbourne.

## Supplementary figure legends

**FIG. S1.**
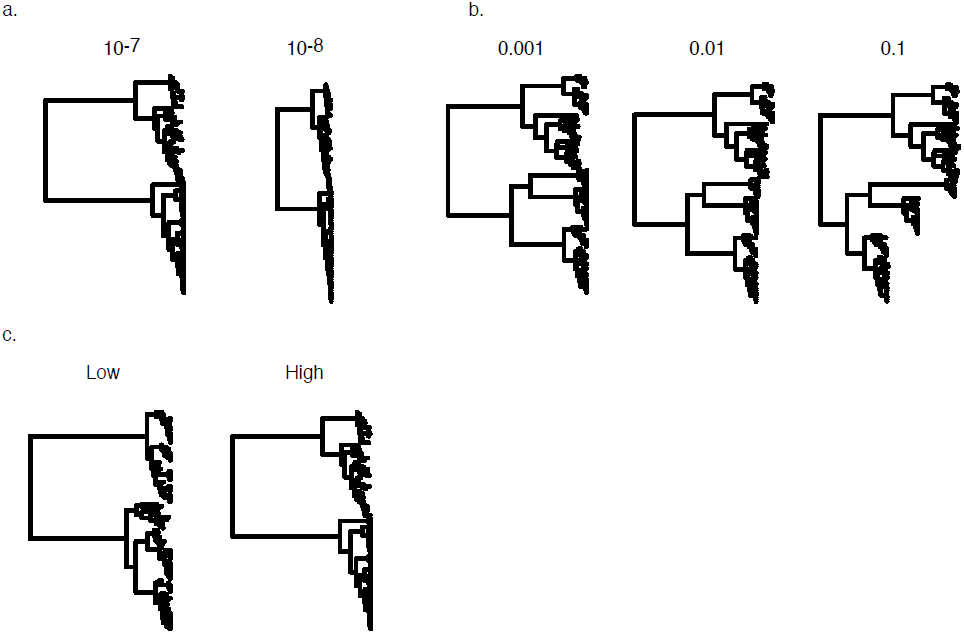
Diagrammatic representations of the different treatments investigated in our simulation study. Simulations of sequence evolution were performed using (*a*) two different substitution rates; (*b*) three levels of among-lineage rate variation; and (*c*) two levels of phylo-temporal clustering.

**FIG. S2.**
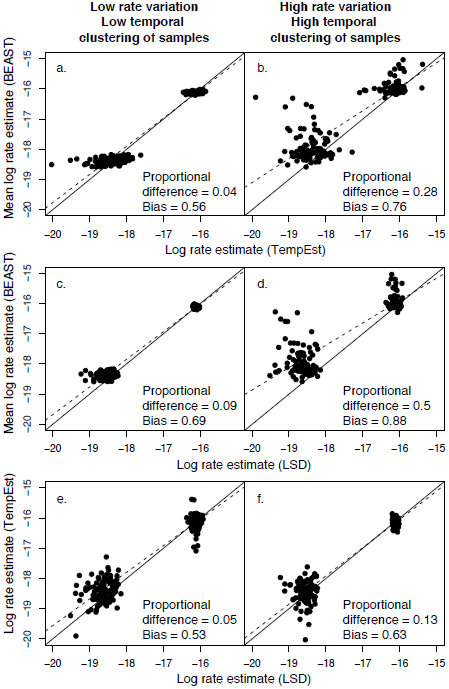
Pairwise comparisons of rate estimates from root-to-tip regression in TempEst, least-square dating in LSD, and Bayesian inference in BEAST. The better the points fit along the solid linear lines, the more congruent one method is with the other. Dashed lines indicate a line of best fit for the estimates. The two distinct clouds of points within each panel represent the high rate and low rate estimates. Proportional difference and bias were calculated as in Duchêne et al. (2016). Proportional difference is the difference in the estimates between two methods, divided by the rate of the first rate estimate ((e_1_ – e_2_) / e_1_). Bias is the proportion of data sets for which the estimate along the x-axis is greater than that along the y-axis.

**FIG. S3.**
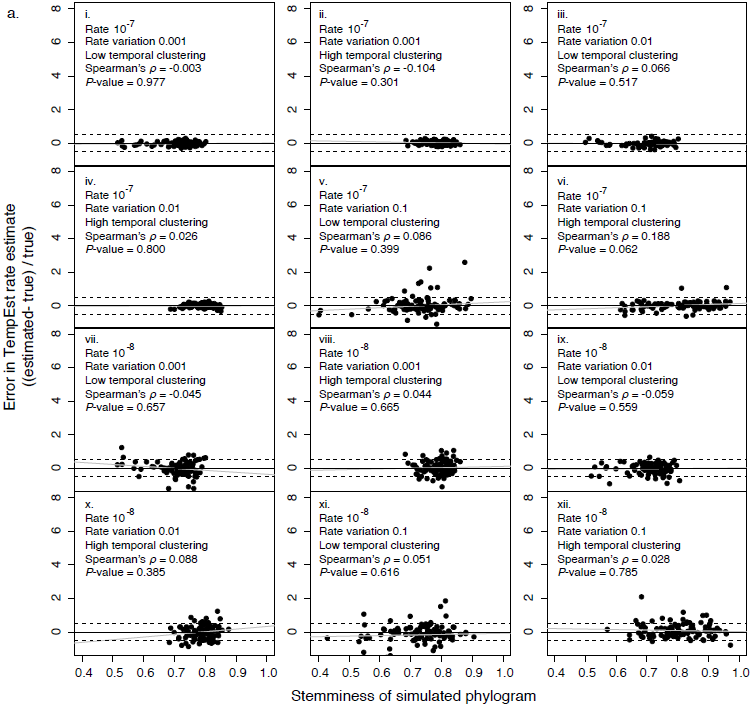
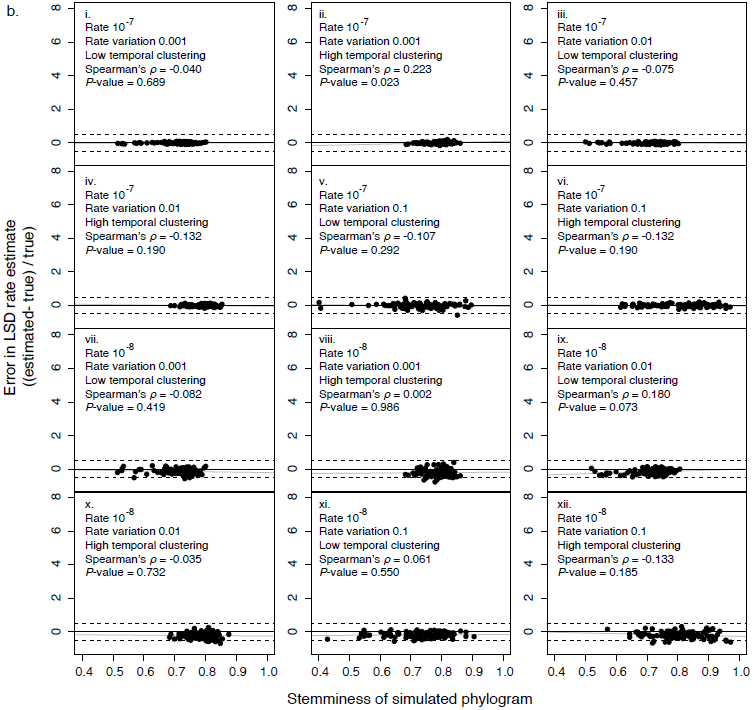
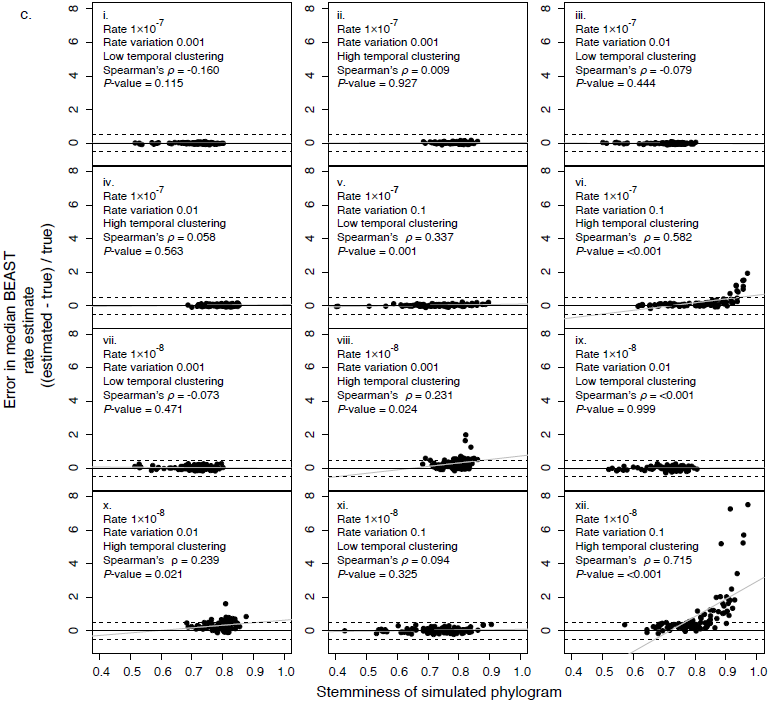
Relationships between phylogenetic stemminess and estimation error for 12 simulation treatments across three methods: (*a*) root-to-tip regression in TempEst, (*b*) least-squares dating in LSD, and (*c*) Bayesian inference in BEAST. Solid dark lines indicate true simulated rates. Dashed lines indicate half a degree of magnitude above or below true rates. Light grey lines indicates lines of best fit for the estimates.

## Supplementary Tables

**Table S1.** Six time-structured mitogenomic data sets analysed in this study.

| Species | Scientific name | Tips | Length<br>(nt) | Age range<br>(years) | Outgroup | Main<br>sources<sup>a</sup> |
| --- | --- | --- | --- | --- | --- | --- |
| Adelie<br>penguin | *Pygoscelis*<br>*adeliae* | 20 | 14,198 | 0–44,000 | *Pygoscelis*<br>*antarctica* | 1 |
| Brown/polar<br>bear | *Ursus arctos* &<br>*U. maritimus* | 32 | 14,609 | 0–122,500 | *Ursus*<br>*americanus* | 2 |
| Dog | *Canis familiaris* | 138 | 14,596 | 0–36,000 | *Canis latrans* | 3 |
| Horse | *Equus caballus* | 167 | 14,910 | 0–42,577 | *Equus asinus* | 4–6 |
| Modern<br>human | *Homo sapiens* | 64 | 14,889 | 0–7,141 | *Homo*<br>*neanderthalens*<br>*is* | 7 |
| Woolly<br>mammoth | *Mammuthus*<br>*primigenius* | 65 | 14,951 | 12,210–<br>46,455 | *Elephas*<br>*maximus* | 8 |

<sup>a</sup> Main publications from which the sequence data were obtained: (1) Subramanian *et al.* (2009); (2) Miller *et al.* (2012); (3) Thalmann *et al.* (2013); (4) Lippold *et al.* (2011); (5) Achilli *et al.* (2012); (6) Orlando *et al.* (2013); (7) Brotherton *et al.* (2013); (8) Gilbert *et al.* (2008).

**Table S2.** Estimates of substitution rates for six time-structured mitogenomic data sets.

| Species | Rate estimate (subs/site/year) |  |  |
| --- | --- | --- | --- |
|  | Root-to-tip regression<sup>a</sup> | Least-squares dating<sup>b</sup> | Bayesian inference<sup>c</sup> |
| Adelie penguin | $3.54 \times 10^{-8}$ | $4.10 \times 10^{-8}$ | $3.37 \times 10^{-8}$ ($1.16 — 5.86 \times 10^{-8}$) |
| Brown/polar bear | $2.72 \times 10^{-8}$ | $2.65 \times 10^{-8}$ | $2.28 \times 10^{-8}$ ($1.5 — 3.22 \times 10^{-8}$) |
| Dog | $1.01 \times 10^{-7}$ | $8.60 \times 10^{-8}$ | $9.32 \times 10^{-8}$ ($7.4 \times 10^{-8} — 1.11 \times 10^{-7}$) |
| Horse | $5.58 \times 10^{-9}$ | $2.26 \times 10^{-8}$ | $3.01 \times 10^{-8}$ ($2.07 — 4.02 \times 10^{-8}$) |
| Modern human | $2.6 \times 10^{-8}$ | $2.1 \times 10^{-8}$ | $2.51 \times 10^{-8}$ ($1.62 — 3.35 \times 10^{-8}$) |
| Woolly mammoth | n/a | $9.59 \times 10^{-8}$ | $1.43 \times 10^{-8}$ ($8.74 \times 10^{-9} — 2.07 \times 10^{-8}$) |

<sup>a</sup> Root-to-tip regression estimates obtained using TempEst (Rambaut et al. 2016); <sup>b</sup> Least-squares dating estimates obtained using LSD 0.3 (To et al. 2016); <sup>c</sup> Bayesian inference estimates obtained using BEAST 1.8.3 (Drummond et al. 2012): median values are provided with 95% credibility intervals detailed within brackets.

